# A survey of DNA methylation polymorphism identifies environmentally responsive co-regulated networks of epigenetic variation in the human genome

**DOI:** 10.1101/189704

**Authors:** Paras Garg, Ricky S. Joshi, Corey Watson, Andrew J. Sharp

**Author notes:** Address for correspondence: Andrew J. Sharp, Department of Genetics and Genomic Sciences, Mount Sinai School of Medicine, Hess Center for Science and Medicine, 1470 Madison Avenue, Room 8-116, Box 1498, New York, NY 10029 USA.

## Abstract

While studies such as the 1000 Genomes Projects have resulted in detailed maps of genetic variation in humans, to date there are few robust maps of epigenetic variation. We defined sites of common epigenetic variation, termed Variably Methylated Regions (VMRs) in five purified cell types. We observed that VMRs occur preferentially at enhancers and 3’ UTRs. While the majority of VMRs have high heritability, a subset of VMRs within the genome show highly correlated variation in *trans*, forming co-regulated networks that have low heritability, differ between cell types and are enriched for specific transcription factor binding sites and biological pathways of functional relevance to each tissue. For example, in T cells we defined a network of 72 co-regulated VMRs enriched for genes with roles in T-cell activation; in fibroblasts a network of 21 coregulated VMRs comprising all four *HOX* gene clusters enriched for control of tissue growth; and in neurons a network of 112 VMRs enriched for roles in learning and memory. By culturing genetically-identical fibroblasts under varying conditions of nutrient deprivation and cell density, we experimentally demonstrate that some VMR networks are responsive to environmental conditions, with methylation levels at these loci changing in a coordinated fashion in *trans* dependent on cellular growth. Intriguingly these environmentally-responsive VMRs showed a strong enrichment for imprinted loci (p<10^−94^), suggesting that these are particularly sensitive to environmental conditions. Our study provides a detailed map of common epigenetic variation in the human genome, showing that both genetic and environmental causes underlie this variation.

## INTRODUCTION

Understanding the causes and consequences of genomic variation among humans is one of the major goals in the field of genetics. Over the past decade, studies such as the Hapmap and 1000 Genomes Projects have resulted in detailed maps of genetic variation in diverse human populations, identifying millions of single nucleotide polymorphisms, copy number variants and other types of sequence variation (The International HapMap Consortium, 2005; The International HapMap Consortium, 2007; The International HapMap Consortium, 2010; Abecasis et al., 2012; Sudmant et al., 2015; Auton et al., 2015). These maps have acted as the catalysts for thousands of genome-wide association studies (Welter et al., 2014), and have provided insights into diverse processes such as mechanisms of human disease, mutation, evolution, migration, selection and recombination (Sabeti et al., 2002; Myers et al., 2005; McEvoy et al., 2011; Zaidi et al., 2013).

However, alterations of the primary DNA sequence are not the only type of genomic variations that occur among humans. In particular there are now well-documented examples of epigenetic marks, such as DNA methylation and histone modifications, that show significant inter-individual variation (Ollikainen et al., 2010; Oey et al., 2015; McDaniell et al., 2010;). However, in contrast to sequence polymorphism, relatively few studies have examined the distribution of epigenetic variation across the genome, and as a result our understanding of the causes and consequences of epigenetic polymorphism remains limited.

Familial and twin studies in human and mice have shown that a substantial fraction of sites showing variable DNA methylation levels are highly heritable, and for some loci this epigenetic polymorphism has been linked with nearby genetic variation (Ollikainen et al., 2010; Oey et al., 2015; Gertz Oey et al., 2011; Grundberg et al., 2012; Grundberg et al., 2013; Gordon et al., 2012; McRae et al., 2014; Busche et al., 2015; Gibbs et al., 2010; Zhang et al., 2010; Gutierrez-Arcelus et al., 2013; Heyn et al., 2013). However, these same studies have also demonstrated that a subset of methylation variation exhibits low heritability (Ollikainen et al., 2010; Grundberg et al., 2012; Grundberg et al., 2013; Gordon et al., 2012; Gervin et al., 2011). While stochastic variation could explain reduced heritability levels, differing environmental exposures such as smoking (Breitling et al., 2011, Joubert et al., 2012; Tsaprouni et al., 2014), diet/in-utero environment (Waterland et al., 2010; Dominguez-Salas et al., 2014; Kok et al., 2015) and stress (Unternaehrer et al., 2012; Donkin et al., 2015) have all been shown to modify the epigenome. In addition, other natural processes such as aging and X chromosome inactivation apparently underlie epigenetic variation of some sites(Christensen et al. 2009; Day et al., 2013; Cotton et al., 2014). Whatever the root cause of epigenetic polymorphism, several studies have demonstrated that a subset of these variations are functionally significant and associate with the expression levels of nearby genes (Gutierrez-Arcelus et al., 2013; Liu et al., 2013). Accordingly there is now substantial interest in elucidating the role of epigenetic variation in a variety of disease phenotypes (Bell et al., 2014; Davies et al., 2014; Huynh et al., 2014; Watson et al., 2016; Multhaup et al., 2015; Benton et al., 2015; Javierre et al., 2010; Lunnon et al., 2014; Pidsley et al., 2014), indicating that the study of epigenetic polymorphism holds significant promise for understanding the molecular etiology of disease.

In this study, we have performed a screen to identify regions of common epigenetic variation using population data derived from five different human cell types. We uncover hundreds of loci in the human genome that exhibit highly polymorphic DNA methylation levels, that we term variably methylated regions (VMRs). We show that VMRs co-localize with other functional genomic features, are enriched for CpGs that influence gene expression, and provide evidence that epigenetic variability at some of these loci is influenced by both genetic and environmental factors. We also show that VMRs form *cis* and *trans* co-regulated networks enriched for transcription factor binding sites and genes with cell-type relevant functions. Finally, consistent with the notion that the epigenome represents a dynamic link between our genome and the environment (Liu et al., 2008; Tammen et al., 2013), we experimentally demonstrate effects of nutrition deprivation on methylation at VMRs in cultured fibroblasts, revealing signatures that overlap those observed in our population-level datasets. Together, our results provide novel insights into the biology of variable methylation across the human genome.

## RESULTS

### Identification of polymorphic DNA methylation in five human cell types

We performed an analysis of inter-individual variation of DNA methylation in five isolated cell types from two human cohorts (Fig. 1A): 1) Primary fibroblasts, EBV-immortalized lymphoblastoid cells, and phytohemagglutinin stimulated primary T cells taken from umbilical cords of 204 newborns (Gutierrez-Arcelus et al., 2013); and 2) sorted glia and neurons from prefrontal cortical tissue from 57 deceased donors (Guintivano et al., 2013). Genome-wide methylation profiles were previously generated for all samples using the Illumina Infinium HumanMethylation450 BeadChip (450k array) (Illumina, San Diego, CA, USA). After filtering (see Methods), we analyzed methylation profiles for 316,452 filtered autosomal CpGs in each of the five cell types. We utilized a sliding window approach (Fig. 1B) to characterize VMRs composed of multiple neighboring CpGs exhibiting consistent polymorphic variation in methylation levels among samples within each cell type.

**Figure 1.**
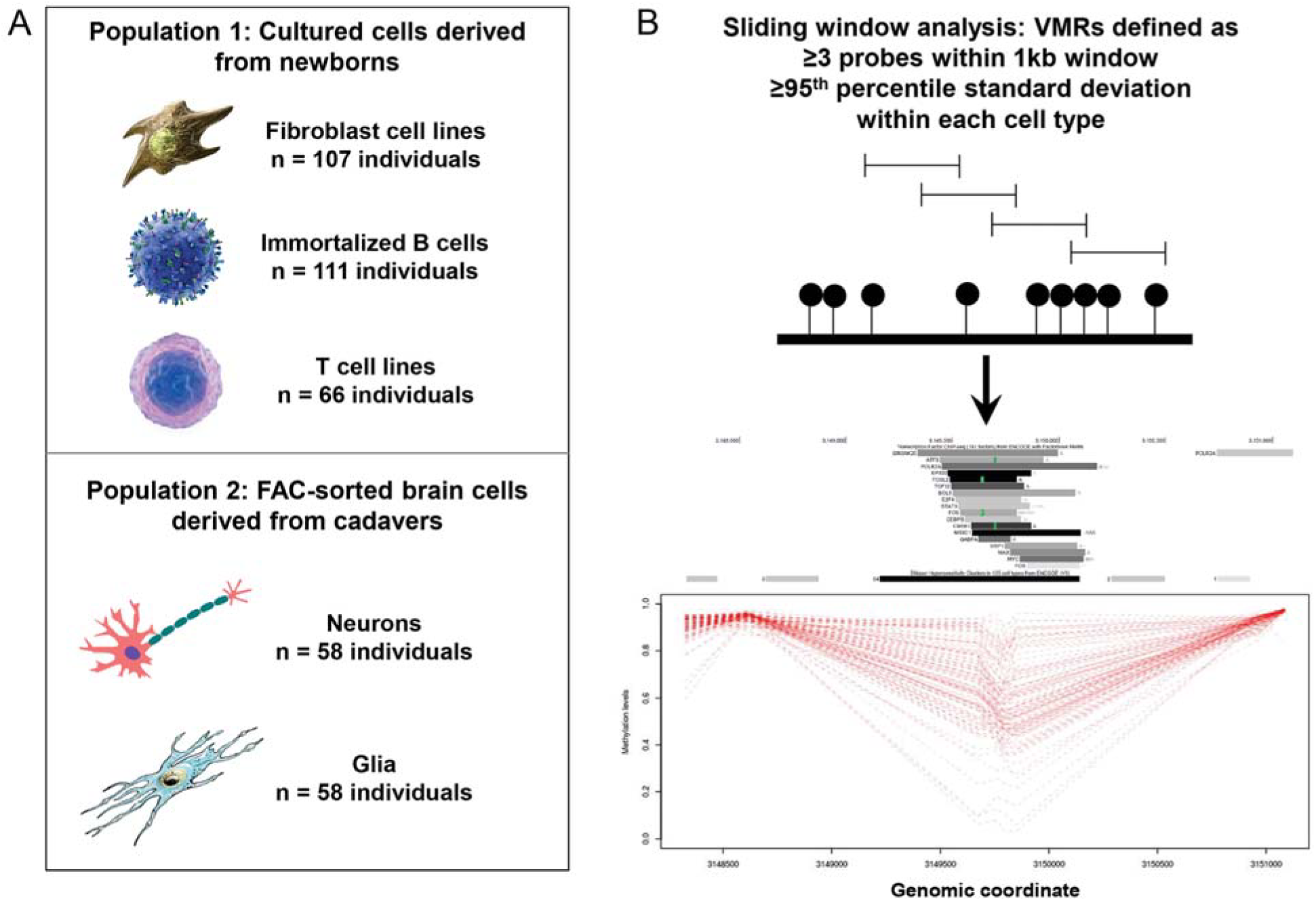
**(A)** We studied population variability of DNA methylation in five different purified cell types derived from blood, skin and brain. **(B)** Utilizing a 1kb sliding window we identified Variably Methylated Regions (VMRs), representing clusters of ≥3 probes within the top 5% of population variability within each cell type. **(C)** An example VMR identified at *PFKP* in fibroblasts. As indicated by the accompanying UCSC Genome Browser tracks, ENCODE data identifies this locus as a DNAseI hypersensitive site and cell-type specific enhancer bound by several different TFs. Dashed red lines represent DNA methylation profiles for each of the 107 cell lines from the GenCord population, showing extreme epigenetic variability at this locus in the normal population.

In total, we identified 537 VMRs in fibroblasts, 1,168 VMRs in T cells, 580 VMRs in B cells, 846 VMRs in neuronal cells and 890 VMRs in glial cells. Hereafter, these VMRs are abbreviated as FVMRs, TVMRs, BVMRs, NVMRs and GVMRs, respectively. Genomic positions and relevant annotations for VMRs partitioned by cell type are provided in Supplementary Table S1. VMRs had a mean size of 875bp, and contained a mean of 6.5 CpGs.

While many characterized VMRs were specific to a given cell type, others were common across cell types and tissues. Examples of cell-type specific and shared VMRs are displayed in Fig. 2A. The extent of VMR sharing between different tissues was related to their relative developmental origin. For example, approximately one third of VMRs identified in glia were also found in neurons, and ~60% of VMRs found in B cells were observed in T cells. In contrast only 22% of VMRs found in fibroblasts were also seen in B cells (Fig. 2B). Between fibroblasts, blood, and brain cells, there were 89 shared VMRs (Fig. 2B). In addition, methylation levels at CpGs within shared VMRs were highly correlated across cell types within tissues, suggesting that observed population variation is plausibly established in precursors of these cell types and maintained, or influenced by common factors and regulatory mechanisms. For example, shared VMRs between T cells and B cells had a mean correlation coefficient of r=0.77 (Supplementary Fig. S1). Likewise for neurons and glia, shared VMR-CpGs were highly correlated (mean r=0.83, Supplementary Figs. S1). However, the same degree of correlation was not observed for comparisons between fibroblasts and T cells (mean r=0.57) or B cells (mean r=0.42, Supplementary Fig. S1).

**Figure 2.**
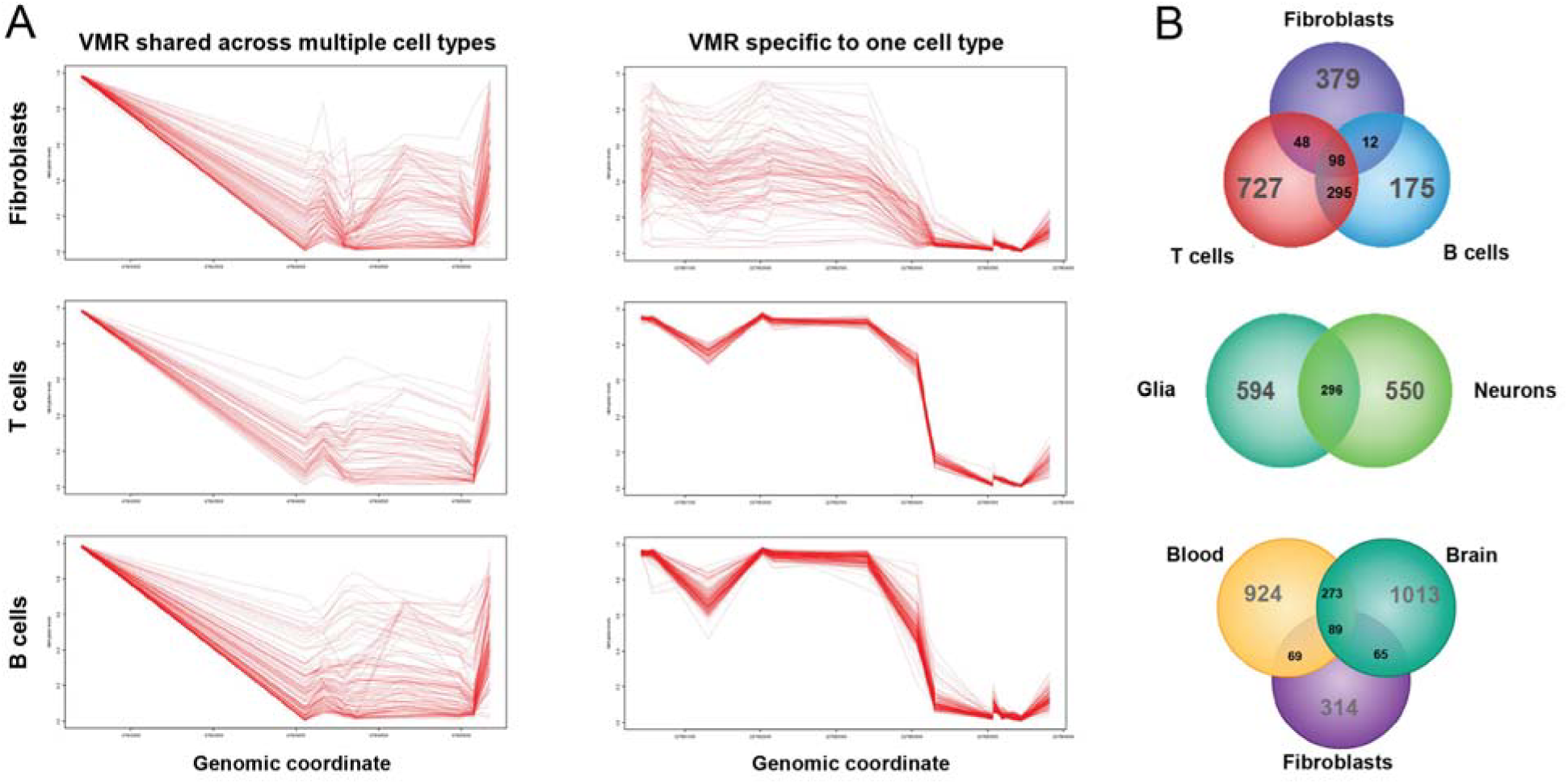
Epigenetic variation in different cell types. **(A)** While some VMRs are common to multiple different cell types, in contrast, other VMRs identified in one cell type show minimal epigenetic variation in other tissues. **(B)** Venn diagram showing the degree of overlap for VMRs found in B-cells, T-cells, Fibroblasts, Neurons and Glia.

### Replication of shared VMRs in a larger cohort

Before extending our analysis of VMRs, we first replicated our method and results in a larger population. We applied our sliding window approach for identifying VMRs to a cohort of 2,680 individuals sampled from the general population (Lehne et al., 2015), identifying 1,312 VMRs. Because this dataset contained methylation profiles generated from peripheral blood, rather than purified cell types, we compared VMRs identified in these controls with shared VMRs that were identified in both B cells and T cells in the Gencord cohort: 339 were also found in the replication cohort, yielding a 32-fold enrichment over that expected by chance (p<2.5×10^−321^).

### VMRs preferentially overlap specific gene/CpG Island features and functional elements in the human genome

Differentially methylated CpGs have been shown to often be enriched in specific regions of the genome and to co-localize with other functional epigenetic signatures (Cedar and Bergman, 2009; Deaton and Bird, 2011; Jones, 2012). In order to gain insight into the genomic context of CpGs in VMRs, we tested the enrichment of these CpGs in relation to various genomic features compared to a background set of CpGs assayed on the array.

We first performed enrichment analysis using Refseq gene and CpG island (CGI) annotations, observing consistent trends across datasets (Supplementary Table S2). Specifically, we noted that in all five of the cell types tested, VMRs were significantly enriched in 3’ UTRs and depleted in 5' UTRs (enrichments ranging from 1.32‐ to 1.73-fold across the different cell types, p=1.4×10^−6^ to p=8.3×10^−18^). Likewise, the depletion of VMRs within 5’ UTRs ranged from 1.36‐ to 1.78-fold (p=1.3×10^−10^ to p=4.0×10^−21^) (Supplementary Table S2). The depletion in 5’ UTRs was also reflected in enrichment tests conducted using CGI annotations, which revealed significant depletions in CGIs and concomitant enrichments in CpG shores, shelves, and sea categories (Supplementary Table S2).

To further explore the co-localization of VMRs with functional genomic regions, we assessed the overlap of FVMRs and BVMRs with Chromatin State Segmentation annotations from a normal human lung fibroblast (NHLF) cell line and an EBV-immortalized lymphoblastoid cell line (GM12878), respectively; these data were previously generated by the ENCODE project (Ernst and Kellis, 2010), and included genome-wide annotations for 15 chromatin states characterized using combined epigenetic signatures from various datasets. Consistent with observed depletions in gene 5’ UTRs and CpG islands, which both tend to occur within or adjacent to gene promoters and transcriptional start sites, we also noted significant depletions of both FVMRs and BVMRs in regions defined by “Active Promoter” chromatin states in respective cell types (Supplementary Table S2). The strongest VMR enrichments in both cell types occurred in chromatin states associated with enhancer activity (Supplementary Table S2).

### VMRs form both cis and trans co-methylated networks that are enriched for genes and transcription factor binding sites with cell-type relevant functions

We next sought to investigate the positional relationships of co-regulated VMRs. In each cell type we constructed pair-wise correlation matrices of all VMRs based on the β-values of the probe with the highest population variance within each VMR. The resulting heat maps of pairwise correlations revealed the presence of strongly co-methylated blocks of CpGs, whose methylation levels varied together in both *cis* and *trans*, and that these patterns were distinct to each cell type (Fig. 3; Supplementary Fig. S2). For example, as shown in Fig. 3A, FVMRs exhibit strong *cis* correlations within several chromosomal regions. Significantly, evidence of strong co-regulation *in trans* can also be seen, with several regions located on multiple different chromosomes also exhibiting strong co-variation in epigenetic state. Visual inspection of the strongest *trans* correlations in fibroblasts located on chromosomes 2, 7, 12 and 17 showed that each of these co-regulated clusters of VMRs corresponded to different members of the *HOX* gene superfamily, suggesting that such VMRs might correspond to coordinately regulated loci with shared biological functions.

**Figure 3.**
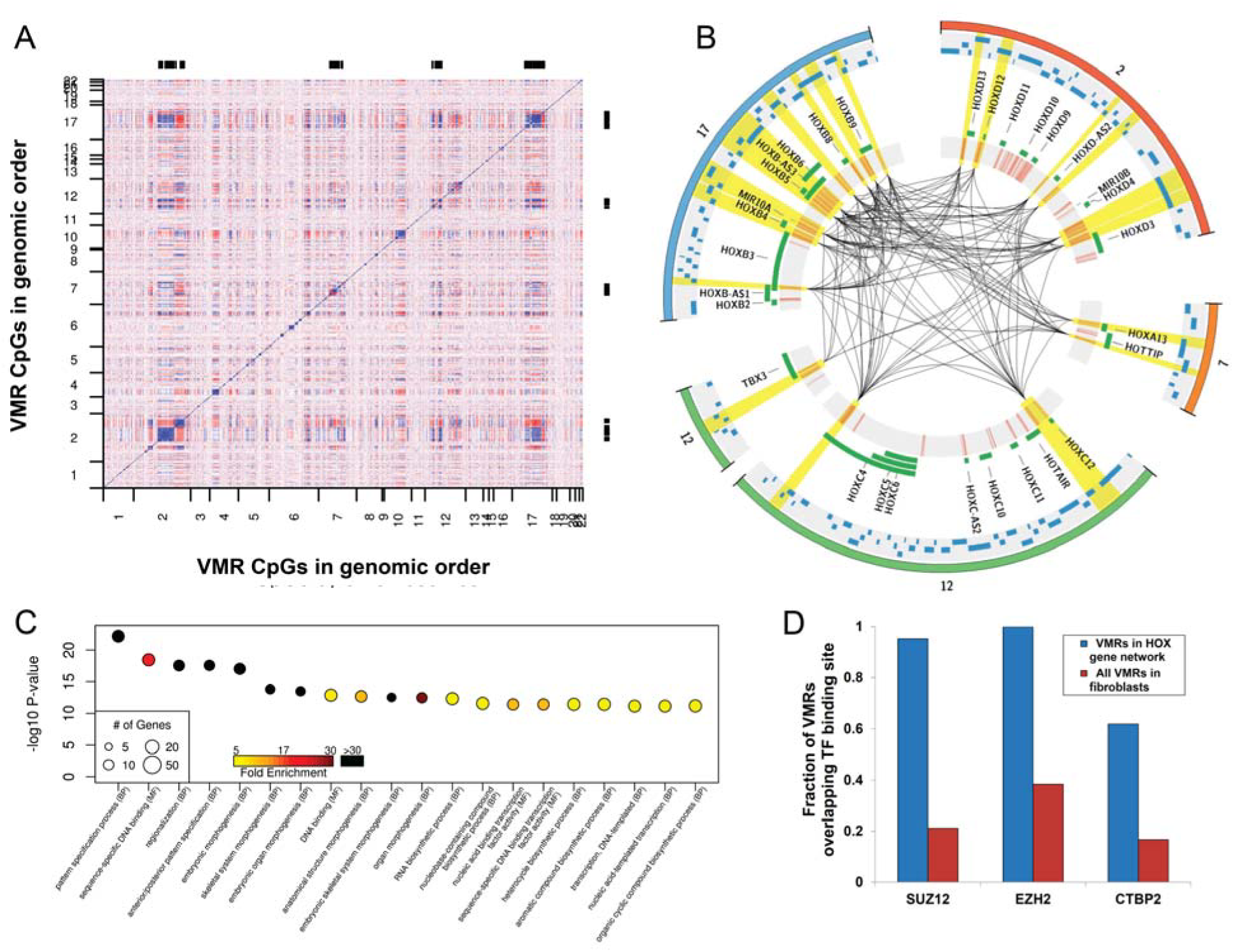
*Cis and trans* co-regulation of VMRs located at functionally related networks of genes that govern key developmental pathways. **(A)** Heat map of pair wise correlation values between all CpGs located within VMRs defined in fibroblasts. CpGs on both axes are ordered by genomic position, revealing the presence of multiple loci located on different chromosomes that show highly correlated methylation levels in *trans*. **(B)** After selecting one CpG per VMR with the highest variance, we applied WCGNA to identify networks of significantly co-regulated VMRs. The Circos plot shows a representation of the largest co-regulated VMR module identified in fibroblasts, which comprises 21 independent VMRs located on four different chromosomes, comprising all four clusters of *HOX* genes, in addition to another developmental regulator *TBX3* (*outer circle*). CpGs within VMRs in the co-regulated module are represented by red tick marks (*inner grey circle*), with black lines joining VMRs that have methylation levels with pair wise absolute correlation values R≥0.7 (*highlighted in yellow*). Green bars show locations of genes at each locus. Blue bars show the location of transcription factor binding sites for SUZ12, EZH2 and CTBP2, all of which are significantly enriched within this co-regulated module. **(C)** Results of Gene Ontology (GO) analysis of genes associated with VMRs in the most significant co-regulated module identified in fibroblasts. We identified highly significant enrichments for multiple biological processes, including body patterning, growth and morphogenesis (Supplementary Table S5). **(D)** Analysis of transcription factor binding sites defined using ChIP-seq (ENCODE Consortium, 2013) showed that VMRs within the co-regulated *HOX* gene module are significantly enriched for SUZ12, EZH2 and CTBP2 binding compared to all VMRs defined in fibroblasts (Bonferroni corrected p=3.8×10^−10^, p=6.0×10^−7^ and p=1.3×10^−3^, respectively). Thus, binding of these TFs represents a potential mechanism by which epigenetic variation could be coordinated at multiple independent loci in *trans*.

Based on this observation, we sought to formally identify signatures of co-regulation among different VMRs. We used weighted gene co-expression network analysis (WGCNA; see Methods)(Zhang and Horvath, 2005; Langfelder and Horvath, 2008), to identify co-methylated networks of VMRs within each cell type. This identified six co-regulated modules in fibroblasts, four in T cells, two in B cells, six in neurons, and two in glia, with each module composed of between 12 and 425 distinct co-regulated VMRs (median module size, n=39) (Supplementary Table S3, Supplementary Fig. S3). Consistent with our initial visual observations, WGCNA identified several co-regulated modules within the set of fibroblast VMRs that included all four human *HOX* gene clusters (Fig. 3B).

In order to assess the biological relevance of these co-regulated VMR networks, we performed Gene Ontology (GO) enrichment analysis on the set of genes linked to the VMRs within each module (Supplementary Tables S4 and S5, Supplementary Fig. S4). Although for many networks the number of associated genes was too small to reach significance at 10% FDR, in four of the five cell types tested we identified enrichments for GO terms that were of direct functional relevance to the specific cell type. The five most significant GO enrichments and associated modules for each cell type are presented in Table 1. For example, in fibroblasts, the most significant functional categories were within the brown module that included multiple *HOX* gene clusters, including terms associated with the basic control of tissue growth and morphogenesis, such as “anterior/posterior pattern specification” (GO:0009952; 82-fold enrichment, FDR q=3.3×10^−16^) and “embryonic morphogenesis” (GO:0048598; 34-fold enrichment, FDR q=8.9×10^−16^). In T cells, the most significant GO enrichments were found for the blue module, made up of 72 co-regulated VMRs enriched for genes involved in T cell function, including the terms “T cell aggregation” (GO:0070489; 15-fold enrichment, FDR q=2.9×10^−6^) and “T cell receptor signaling pathway” (GO:0050852; 15-fold enrichment, FDR q=7.4×10^−5^). In glial cells, significantly enriched terms included a module consisting of 425 VMRs linked to genes associated with “negative regulation of neurogenesis” (GO:0050768; 4.6-fold enrichment, FDR q=1.2×10^−5^). Finally, in neurons, the most strongly associated functional categories were with a module comprised of 112 VMRs including the GO term “negative regulation of synaptic transmission” (GO:0050805; 13-fold enrichment, FDR q=0.09). Complete lists of enriched GO terms and modules are provided in Supplementary Table S5.

**Table 1.** The top three Gene Ontology terms associated with co-regulated VMR modules found in each cell type.

| Cell Type | GO Term ID | Gene Ontology (GO) Term | Enrichment | FDR |
| --- | --- | --- | --- | --- |
| **Fibroblasts** | GO:0007389 | Pattern specification process | 42.7 | 3.07E-20 |
|  | GO:0009952 | Anterior/posterior pattern specification | 81.8 | 3.32E-16 |
|  | GO:0003002 | Regionalization | 54.2 | 3.32E-16 |
| **T Cells** | GO:0007159 | Leukocyte cell-cell adhesion | 14.2 | 2.94E-06 |
|  | GO:0042110 | T cell activation | 15.3 | 2.94E-06 |
|  | GO:0070489 | T cell aggregation | 15.3 | 2.94E-06 |
| **B Cells** | GO:0002761 | Regulation of myeloid leukocyte differentiation | 11.8 | 0.1496 |
|  | GO:1902105 | Regulation of leukocyte differentiation | 7.2 | 0.1658 |
|  | GO:0070233 | Negative regulation of T cell apoptotic process | 50.0 | 0.1658 |
| **Glia** | GO:0050768 | Negative regulation of neurogenesis | 4.6 | 1.18E-05 |
|  | GO:0051961 | Negative regulation of nervous system development | 4.2 | 3.33E-05 |
|  | GO:0045665 | Negative regulation of neuron differentiation | 4.6 | 0.0002 |
| **Neuron** | GO:0007611 | Learning or memory | 6.2 | 0.0905 |
|  | GO:0050805 | Negative regulation of synaptic transmission | 13.2 | 0.0905 |
|  | GO:1901019 | Regulation of calcium ion transmembrane transporter activity | 13.0 | 0.0905 |

Based on the *trans* nature of these co-regulated VMR networks, we hypothesized that coordinated epigenetic regulation of these sites might be based on the binding of specific trans-acting factors to the members of each VMR network. We therefore analyzed the overlap of each VMR WGCNA module with validated transcription factor binding sites (TFBS) for 161 different transcription factors (TFs) studied by the ENCODE project (ENCODE Consortium, 2013). We observed significant enrichments for TFBS in several VMR modules that were specific to each cell type (Supplementary Table S6). The top three enriched TFBS per cell type are provided in Table 2. In several instances, the most significant TFBS enrichments converged on modules highlighted by GO analyses. For example, EBF1 and RUNX3, which are both involved in lymphocyte differentiation and proliferation (Heltemes-Harris and Farrar, 2012), were the most significantly enriched TFs in the blue module in T cells (RUNX3, 2.4-fold enrichment, p=5.3×10^−8^; EBF1, 2.6 fold enrichment, p=7×10^−8^). Similarly, in fibroblasts, TFBS for SUZ12 (3.5-fold enrichment, Fisher’s. p=3.4×10^−10^) and EZH2 (2.4-fold enrichment, p=8.0×10^−9^), were the most significantly enriched among VMRs of the module that included multiple HOX-genes (Fig. 3D). Prior studies have shown that as part of the polycomb complex, SUZ12 and EZH2 have roles in the establishment of epigenetic modifications, and specifically in the regulation of *HOX* genes (Cao et al, 2008).

**Table 2.**
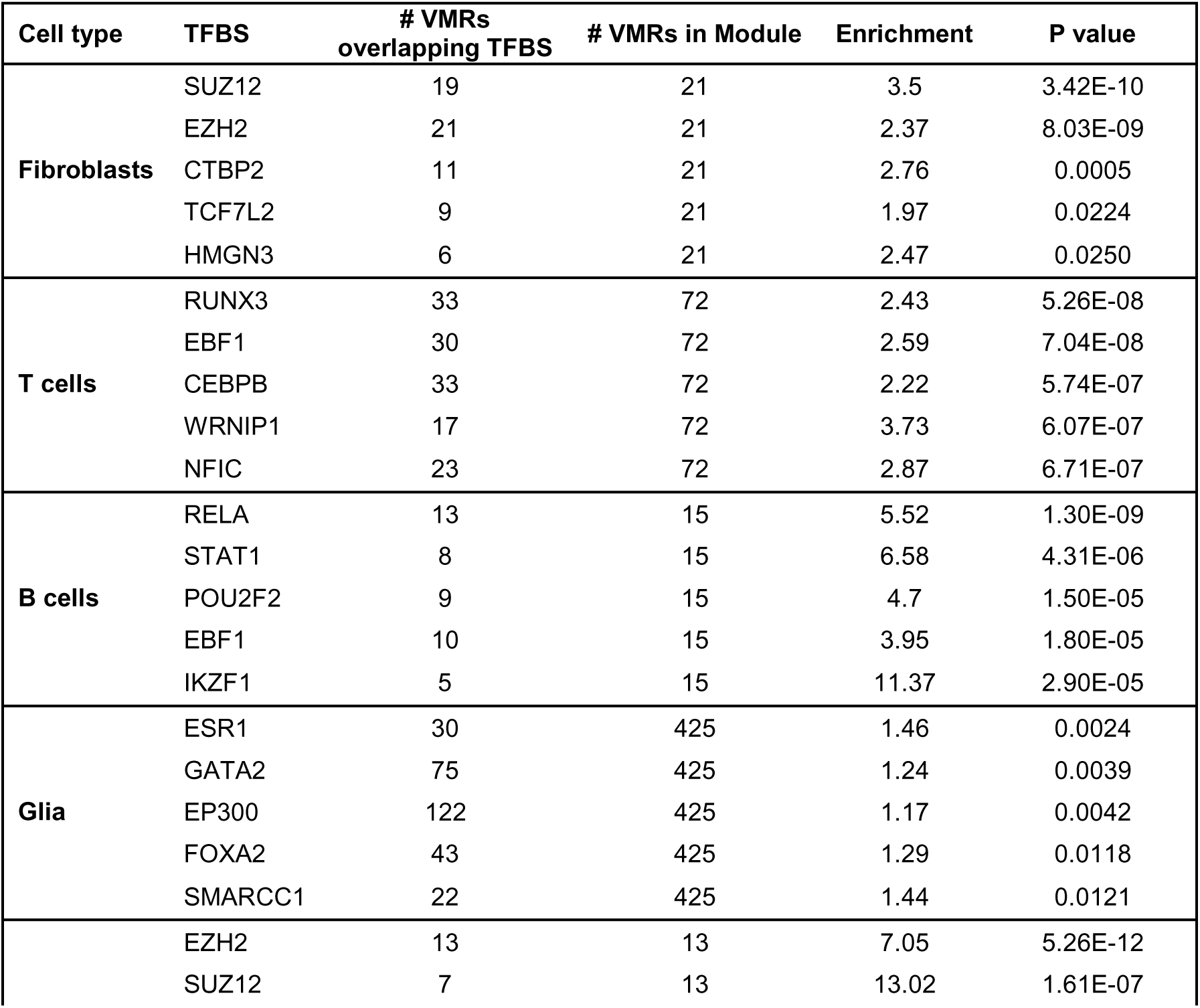
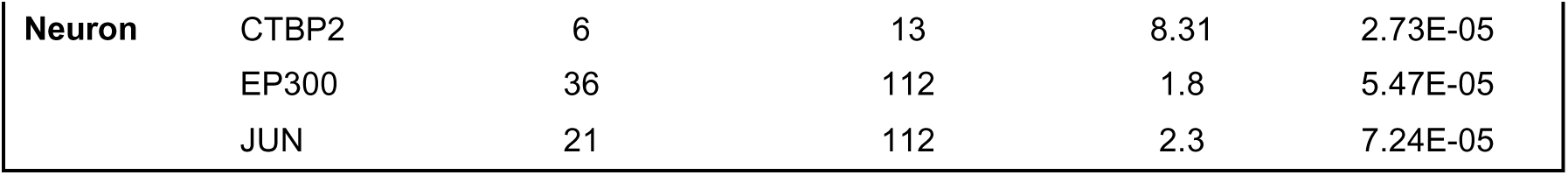
Top 5 Transcription Factor Binding Sites overlapping with VMRs in various WGCNA modules in 5 cell types.

### Methylation levels at VMRs are influenced by both heritable and non-heritable factors

Motivated by the signatures of co-methylation observed in our VMRs, we next sought to broadly explore the potential underlying factors associated with the regulation of VMR methylation variability. To do this, we first assessed the relationships between CpGs within VMRs, genetic variation, and gene expression. We tested for enrichment of FVMRs, BVMRs and TVMRs with previously described CpG methylation:gene expression associations (eQTMs) and CpG methylation:SNP associations (*cis* mQTLs) in fibroblasts, B cells and T cells (Gutierrez-Arcelus et al., 2013). We observed significant enrichments for VMRs in all three cell types for both CpGs that function as eQTMs and those linked with mQTLs, with enrichments of 18.8-, 3.2-, and 3.6-fold in eQTMs, and 4.4-, 6.5-, and 5.1-fold for association with mQTLs in FVMRs, BVMRs, and TVMRs, respectively (all p-values <10^−53^, Supplementary Table S2). Similarly, 174 (32.4%) of FVMRs, 796 (68.1%) of TVMRs and 343 (59.1%) of BVMRs overlapped with mQTLs defined in prior analyses (Gutierrez-Arcelus et al., 2013). To further investigate the relationship of VMRs with underlying genetic variation we used methylation heritability estimates characterized in peripheral blood leukocytes from a cohort of 117 families (McRae et al., 2014). Overlaying heritability estimates onto VMR-CpGs across the five cell types revealed that methylation levels for CpGs within VMRs showed significantly increased heritability compared to non-VMR CpGs (Fig. 4A). Thus, epigenetic variation at VMRs is often associated with nearby gene expression, and methylation levels at many VMRs shows strong evidence of being under local genetic control.

**Figure 4.**
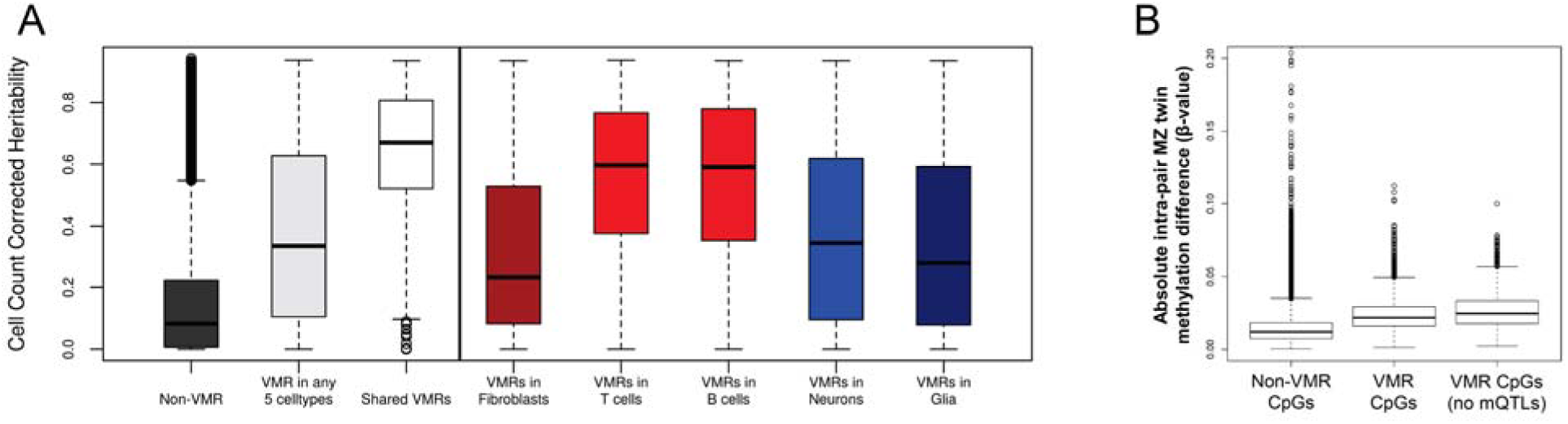
Methylation levels at VMRs are influenced by heritable and non-heritable factors. **(A)** Cell count corrected heritability (Grundberg et al., 2013) for VMRs in five cell types. VMRs found in >1 cell type show significantly higher heritability, suggesting these are mostly under genetic control **(B)** Methylation differences found within 13 pairs of monozygotic twins. CpGs that lie within VMRs show significantly increased MZ twin divergence compared to other CpGs, suggesting an environmental influence on methylation levels at VMRs. This effect is even more pronounced after excluding those VMRs known to be under genetic control.

However, despite this evidence for genetic influences underlying a large fraction of epigenetic variability, the existence of co-regulated modules of VMRs *in trans* led us to hypothesize that a subset of epigenetic variation might be linked to non-genetic influences, such as differing environmental exposures. To further explore the influence of non-heritable factors on the epigenetic state of VMRs, we analyzed methylation profiles from 13 monozygotic (MZ) twin pairs (McRae et al., 2014). Previous studies have shown increased discordance of DNA methylation levels between MZ twins with age, presumably due to differing environmental exposures and/or stochastic processes (Fraga et al., 2005). We first identified a total of 1,411 VMRs (9,079 CpGs) in these twins (Supplementary Table S7). Based on the premise that epigenetic differences between MZ twin pairs provides a measure of the non-genetic component of epigenetic variability, at each CpG we calculated the mean absolute methylation discordance for all autosomal CpGs within each MZ twin pair. We observed a highly significant increase in MZ twin discordance for CpGs within VMRs versus non-VMR CpGs (p<10^−300^) (Fig. 4B). Furthermore, after removing those CpGs known to be associated with mQTLs, *i.e.* those under genetic control, the degree of MZ twin discordance in the remaining VMRs increased substantially (Fig. 4B). These observations provide strong support for the influence of environmental effects on methylation variability at a subset of VMRs.

### Experimental evidence for environmental influences on DNA methylation from a cell culture model

To experimentally verify whether methylation levels at some VMRs are responsive to environmental cues, we performed cell culture experiments in which we grew genetically identical fibroblasts under different environmental conditions, varying levels of nutrient deprivation and cell density with time (Fig. 5A). Skin fibroblasts from a single normal male (GM05420) were seeded in parallel from a single master culture into eight separate flasks, and allowed to grow under normal or low-nutrient conditions, achieving varying levels of cell density at each time point. Every 48 hours one flask was harvested from each nutrient regime, DNA extracted and profiled on the 450k array, resulting in DNA methylation profiles for nine samples (see Methods).

**Figure 5.**
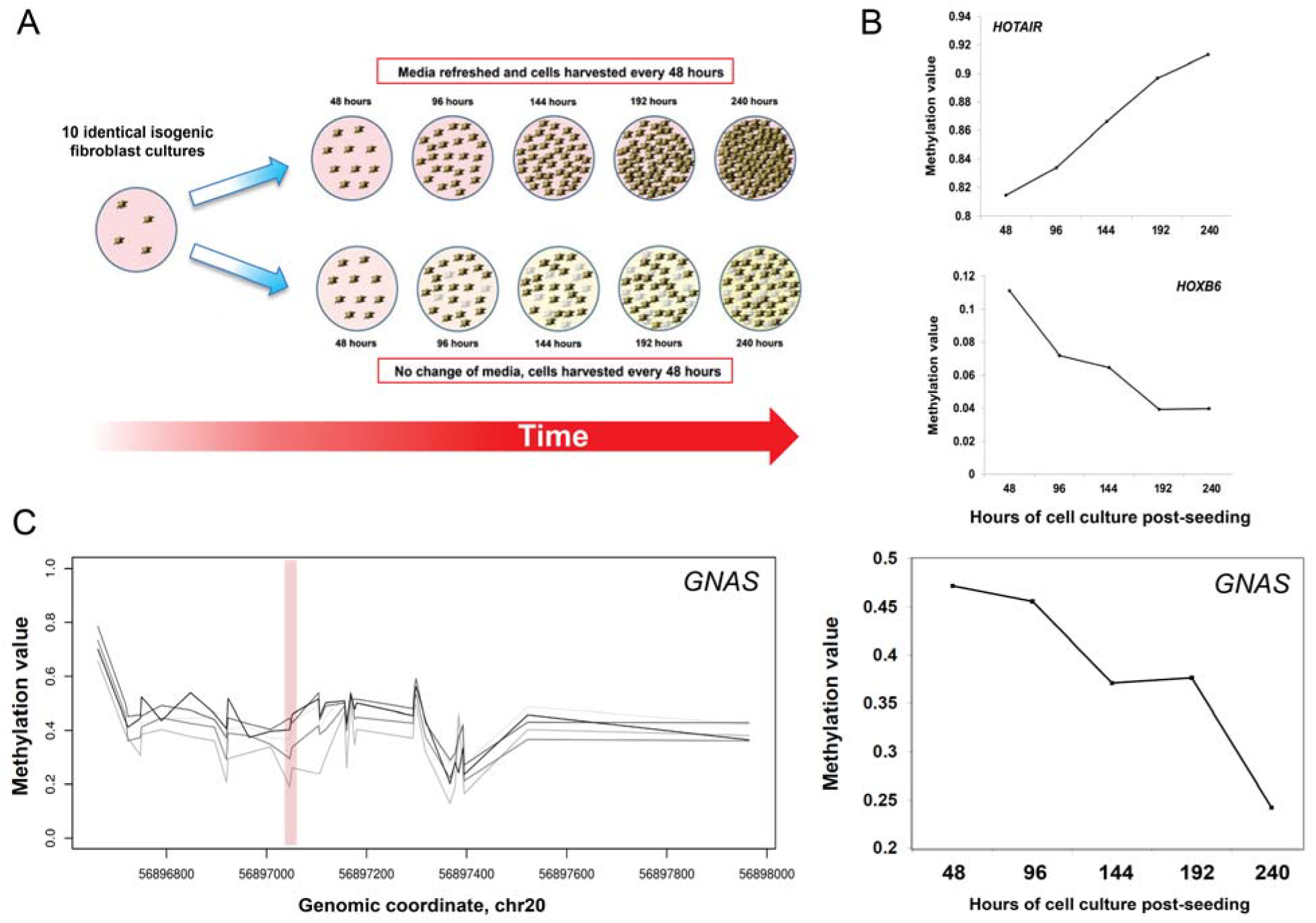
Experimental manipulation of DNA methylation using cell culture shows enrichment for VMRs at *HOX* genes and imprinted loci. **(A)** To directly assess the effect of varying environmental conditions on epigenetic state, we grew genetically-identical fibroblasts under conditions of increasing cell density and nutrient deprivation. Cells from a single human fibroblast line were seeded in parallel at low density in ten culture flasks, and allowed to grow continuously for up to 10 days, either with or without regular change of media. Every 48 hours one flask was harvested and genome-wide DNA methylation patterns profiled. **(B)** Applying a sliding window approach identified 142 VMRs where methylation levels showed robust changes with varying culture conditions, including loci at several *HOX* genes and multiple imprinted loci. Gene ontology analysis of VMRs induced by cell culture showed enrichments for fundamental control of growth, including similar GO categories to the co-methylated network identified in fibroblasts from the Gencord cohort (data not shown). **(C)** Environmentally-responsive VMRs induced by cell culture showed a 47-fold enrichment for probes within the differentially methylated regions associated with nine different imprinted genes (p=1.5×10^−94^). The left plot shows methylation profiles at the imprinted region of *GNAS*, which was also identified as a VMR in cultured fibroblasts. Each line shows the methylation profile at a different time point, with lighter shades of grey with increasing time. The right plot shows the change in methylation level with time at a single CpG (cg22407822) within the *GNAS* VMR.

We applied the same sliding-window method to identify VMRs in these cultured isogenic fibroblasts, identifying 162 putatively “environmentally responsive” VMRs. This included many of the same VMRs identified previously in our population-based analysis of umbilical cord-derived fibroblasts (Supplementary Table S8), with a 7.3-fold enrichment for overlap between these two sets of VMRs (p=1.4×10^−47^). Examples of VMRs showing changes in methylation level with culture conditions are shown in Fig. 5B.

Concordant with our population analysis, GO analysis of the 162 VMRs from cultured isogenic fibroblasts revealed enrichments for *HOX* genes, as well as the several of the same GO terms associated with the co-regulated FVMR modules (Supplementary Table S9). Strikingly, these environmentally responsive VMRs were also enriched 47-fold for CpGs within known imprinted loci versus the null (P=1.5×10^−94^). This included overlaps with differentially methylated regions associated with the imprinted genes *PPIEL*, *MKRN3*, *MAGEL2*, *SNRPN*, *PEG3*, *L3MBTL1*, *MEST*, *PLAGL1*, and *GNAS* (Fig. 5C) (Court et al., 2014).

## DISCUSSION

Here we surveyed variation in DNA methylation patterns in five purified human cell types, identifying hundreds of genomic loci that exhibit a high degree of epigenetic polymorphism in the human population: we term these ‘Variably Methylated Regions’ or VMRs. We observed that VMRs are enriched for various functional genomic features, most notably enhancers, suggesting a potential role in regulating gene expression patterns. Unexpectedly, we found that many VMRs form co-regulated networks both *in cis* and *in trans*, with multiple VMRs spread across different chromosomes at which methylation levels vary in a coordinated fashion. These co-regulated networks were specific to each cell type, had reduced heritability, and were also enriched for gene sets with cell-type relevant functions. For example, we observed VMR networks associated with genes enriched for learning/memory and synaptic transmission in neurons, regulation of nervous system development in glia, and T cell activation in T cells. These observations suggest that some VMRs represent loci that form co-regulated pathways that are implicated in the regulation of genes with cell-type specific functions. The dispersed nature of these co-regulated VMR networks indicates that they are potentially regulated by *trans* acting factors, and consistent with this we found significant enrichments for relevant transcription factor binding sites associated with some networks.

While many VMRs are influenced by local genotypes, our analyses of monozygotic twins and *in-vitro* culture of genetically identical fibroblasts cell lines clearly demonstrates that epigenetic variation at some VMRs is linked to environmental factors. Indeed, using isogenic fibroblast cultures derived from a single individual that were grown under different environmental conditions, we were able to replicate many of the same VMRs found in our original population analysis, thus showing definitively that epigenetic variation at these loci is an environmentally inducible trait. Intriguingly these environmentally-responsive VMRs showed a strong enrichment for imprinted loci (p<10^−94^), suggesting that these genes are particularly sensitive to environmental conditions. This observation that varying cell culture conditions result in epigenetic alterations across the genome, presumably accompanied by changes in gene expression, highlights that the use of cultured cells for investigating epigenetic phenomena should be approached with caution. We suggest that unless carefully controlled, variations in cell culture conditions could easily introduce significant epigenetic and transcriptional changes that could confound many *in vitro* studies.

VMRs in fibroblasts comprised co-regulated modules that included all four *HOX* gene clusters that are each located on different chromosomes. Using validated transcription factor binding sites, we found a significant enrichment for transcription factors EZH2 and SUZ12 at these VMR sites associated with *HOX* genes. These two transcription factors are components of the Polycomb Repressive Complex 2 (PRC2), which functions as a histone H3K27-specific methyltransferase and regulates both epigenetics and expression of *HOX* genes (Cao et al., 2008). Thus, we propose a model where coordinated variation of DNA methylation at multiple loci in *trans*, corresponding to a network of co-regulated genes, is under the control of transcription factor binding in response to physiological and/or environmental cues. In the case of the *HOX* gene network in cultured fibroblast cell lines, such cues could be the availability of nutrients, local cell density and other growth conditions, allowing the cells to modify their growth trajectories in response to the prevailing environment. Consistent with this model, recent observations were made in macrophages, a type of immune cell that has a variety of roles in different tissues around the body, which mirror our findings. Two prior studies showed that the epigenetic state of enhancer elements in these cells responds to the tissue microenvironment in which they reside, and is regulated by networks of tissue‐ and lineage-specific transcription factors that drive divergent programs of gene expression (Gosselin et al., 2014; Lavin et al., 2014). Studies of chromatin accessibility have also shown that manipulating the presence of specific transcription factors can lead to global modification of epigenetic state at multiple loci *in trans* (Buenrostro et al., 2015).

One of the strengths of this study is that we specifically utilized purified cell types for our analysis, some of which were also of homogeneous age. This has the advantage of removing the confounder of both cellular heterogeneity and age effects, both of which are known to influence DNA methylation (Christensen et al. 2009; Day et al., 2013; Houseman et al., 2015). Such differences would otherwise result in many false positive VMRs due to underlying differences in cell fractions or age among individuals. Furthermore, we also utilized a window-based approach for defining VMRs. This requirement for the presence of multiple neighboring CpGs with high population variance means our results should be robust to potential effects of underlying sequence variants, which can artifactually influence reported methylation levels at single probes.

One of the limitations of this analysis is that we used methylation profiles from the Illumina 450k array, which targets only a small subset (~3%) of CpGs in the human genome, and has coverage that is biased towards gene promoters and CpG islands. As such, the maps of VMRs we provide here are far from comprehensive, and future work that utilizes more comprehensive approaches *(e.g.* whole genome bisulfite sequencing) will undoubtedly provide more complete genomic maps of epigenetic variation.

However, to our knowledge currently no such datasets on a population-scale are available. One other potential caveat is that the methylation profiles for B cells, fibroblasts and T cells were all generated from cells that had been cultured *in vitro*, and furthermore the B cells were also immortalized by Epstein-Barr virus infection, a process which is known to induce widespread epigenetic changes (Grafodatskaya et al., 2010). However, we observed good replication of the VMRs identified from cultured/immortalized B cells and T cells in an independent cohort where DNA was extracted from uncultured blood, indicating that many of these same VMRs observed even in immortalized B cells are also present natively.

In conclusion our study of DNA methylation polymorphism provides novel insights into the nature and function of epigenetic variation. The coordinated response we observed where methylation levels at networks of multiple genomic regions varies in response to the local environment is consistent with popular theories that the epigenome can indeed act as an interface between the genome and environment (Liu et al., 2008; Tammen et al.,2013, Bell and Beck, 2010).

## METHODS

### Data processing and statistical analysis

We obtained DNA methylation data generated using the Illumina 450k HumanMethylation BeadChip from two published studies. We utilized data from the Gencord cohort from the EMBL-EBI European Genome-Phenome Archive (https://www.ebi.ac.uk/ega/) under accession number EGAS00001000446, representing 107 fibroblast cultures, 66 T-cell cultures and 111 immortalized B-cell cultures derived from a cohort of newborns (Gutierrez-Arcelus et al., 2013). We also utilized methylation data representing FAC-sorted glial and neuronal cells from 58 deceased donors downloaded from GEO (http://www.ncbi.nlm.nih.gov/geo/) under accession number GSE41826 (Guintivano et al., 2013). Prior to analysis for methylation variation, each dataset underwent several filtering and normalization steps, as follows. In each individual, probes with a detection p>0.01(mean n=348 per sample) or mapping to the X or Y chromosomes were removed. 482,421 probe sequences (50-mer oligonucleotides) were remapped to the reference human genome hg18 (NCBI36) using BSMAP, allowing up to 2 mismatches and 3 gaps, retaining those 470,681 autosomal probes with unique genomic matches. Probe coordinates were converted to hg19 using *liftover*. Probes that overlapped SNPs identified by the 1000 Genomes Project (minor allele frequency ≥0.05) either including or within 5bp upstream of the targeted CpG (n=9,409 autosomal probes) were discarded, as such variants can introduce biases in probe performance. We also removed probes overlapping copy number variants of ≥5% frequency in CEU HapMap samples (Conrad et al., 2010). One pair of neuronal/glial samples was excluded on the basis that they showed discrepant gender, as determined by PCA analysis of β-values on the sex chromosomes. For sliding window analysis, we sub-selected 316,452 autosomal 1kb windows containing 3 or more probes. CpGs targeted by these probes were then annotated based on their position relative to RefSeq genes using BEDTools v2.17.

### Variably Methylated Regions

To identify regions of common highly variable methylation that should be robust to fluctuations in single probes, we chose an approach to identify loci containing multiple independent probes showing high population variance. For each probe, we calculated the standard deviation (SD) of the β-value across all individuals in each cell type. We then utilized a 1kb sliding window based on the start coordinate of each probe, beginning at the most proximal probe on each chromosome and moving down consecutively to the last probe on each chromosome. We defined VMRs as those 1kb regions containing at least 3 probes ≥95^th^ percentile of SD in that cell type, with an additional criterion that at least 50% of the probes in that window were also ≥95^th^ percentile of SD.

### Network Analysis and Gene Ontology Analysis

To identify potential co-regulation relationships among VMRs, we applied Weighted Gene Correlation Network Analysis (WGCNA) to each set of VMRs identified per cell type (Langfelder and Horvath, 2008). Input values for each VMR were first reduced to a single data point per individual by averaging β-values for the multiple probes within each VMR. We generated adjacency matrices by raising the correlation matrix to the power of 6, which was then transformed into topological overlap matrix (TOM). VMRs were then classified into modules using hybrid dynamic tree cutting with a minimum cluster size of 10 probes. VMRs in each module were selected at Module Membership value ≥0.7. We associated VMRs with gene annotations based on either their localization within ±2kb of Refseq transcription start sites, overlap with DNAseI hypersensitive sites that showed significant association *in cis* with gene expression levels within ENCODE cell lines (Sheffield et al., 2013), and significant associations between methylation and gene expression levels (eQTMs) in T cells, B cells, fibroblasts (Gutierrez-Arcelus et al., 2013). For each module with at least 10 associated genes, we performed Gene Ontology enrichment analysis using GOrilla (http://cbl-gorilla.cs.technion.ac.il/)(Eden et al., 2009).

### Enrichment analysis of transcription factor binding sites

We downloaded the track of Uniform transcription factor binding sites (TFBS) from the UCSC Genome browser, containing experimentally determined binding sites for 162 transcription factors. As the precise boundaries of some VMRs were not well defined, we extended TFBS coordinates by ±500bp prior to overlap with the set of VMRs identified in each cell type. Enrichment analysis for TFBS to occur within each module of co-regulated VMRs identified by WCGNA versus the background was performed using a Fisher's exact test. In each cell type, only TFBSs with ≥10 overlaps with VMRs were considered, and we applied a Bonferroni correction to p-values based on the total number of TFs tested (n=162).

### Fibroblast cell culture and methylation profiling

A growing culture of human skin fibroblasts from a normal male individual (GM05420) was obtained from Coriell Institute for Medical Research (Camden, NJ). Cells were grown in RPMI1640 media supplemented with 1mM L-glutamine, 10% FBS and 100u/L each of penicillin and streptomycin. A single vial of fibroblasts was initially grown in a 2ml culture plate, with media changed every 24 hours. Once the cells attained 80% confluency they were trypsinized and split equally into two T25 flasks. Each flask was treated identically, with media changed every 24 hours until the cells achieved 80% confluency (approximately 7 days after seeding). Both cultures were then trypsinized, mixed, and the cells seeded equally into a total of nine T25 flasks, which were then harvested at set time points (TP) under different culture regimes, as follows:

1. Harvested immediately
2. Time Point (TP) 1 - harvested after 48 hours
3. TP2 - fresh media given at TP1 and then harvested after a further 48 hours
4. TP3 - fresh media given at TP1 and TP2, and then harvested after a further 48 hours
5. TP4 - fresh media given at TP1, TP2 and TP3, and then harvested after a further 48 hours
6. TP5 - fresh media given at TP1, TP2, TP3 and TP4, and then harvested after a further 48 hours
7. TP2a – No change of initial media, harvested after 96 hours
8. TP4a – No change of initial media, harvested after 192 hours
9. TP5a – No change of initial media and then harvested after 240 hours

At each time point, cells were harvested by trypsinization, pelleted by centrifugation, and frozen at −20 Celsius. Once all cultures were harvested, DNA was extracted in a single batch using the Qiagen DNeasy blood and tissue kit and these samples processed together on a single chip using the Illumina 450k HumanMethylation BeadChip according to manufacturer’s instructions. The resulting data were then processed and normalized as described above, and VMRs across these nine samples defined as 1kb regions containing at least 3 probes ≥95^th^ percentile of SD, with an additional criterion that at least 50% of the probes in that window were also ≥95^th^ percentile of SD.

## DATA ACCESS

Methylation array data from cell line GM05420 have been deposited in the NCBI Gene Expression Omnibus (GEO) (http://www.ncbi.nlm.nih.gov/geo) under accession number GSE76836.

## AUTHORS’ CONTRIBUTION

PG, RJ and AJS designed the research. PG performed the bioinformatics analysis. RJ prepared the biological material and CW performed data analysis for time-point experiments. PG, CW and AJS wrote the manuscript with valuable contribution from RJ. AJS coordinated the study. All authors read and approved the final manuscript.

## ACKNOWLEDGMENTS

This work was supported by NIH grant HG006696 and research grant 6-FY13-92 from the March of Dimes to AJS, and a Beatriu de Pinos Postdoctoral Fellowship to RSJ (2011BP-A00515).

---

**Supplementary Fig. S1. Pairwise correlation between shared VMRs reveals varying levels of similarity across cell types**. VMRs found in fibroblasts show relatively low correlations with other cells types, whereas there is much greater similarity in VMRs between T-cells and B-cells (both of which are types of blood cell), and even greater similarity between VMRs found in glia and neurons (both of which are derived from brain).

**Supplementary Fig. S2. Heat maps showing pair wise correlation values between all CpGs located within VMRs defined in neurons, glia, B cells and T cells**. In each plot, CpGs on both axes are ordered by genomic position, revealing the presence of multiple loci located on different chromosomes that show highly correlated methylation levels in *trans*.

**Supplementary Fig. S3. Examples of networks of genes associated with co-regulated VMRs identified in four cell types**. The outermost circle in each Circos plot represents segments of each chromosome. Gene names are shown inside. Blue tick marks on the inner grey shaded band represent each VMR. The black curved lines connect the VMRs in the network that have methylation levels with pair wise absolute correlation values R≥0.7.

**Supplementary Fig. S4. Results of Gene Ontology (GO) analysis of genes associated with networks of co-regulated VMRs identified by WCGNA. (A)** T cells, **(B)** glia, and **(C)** neurons.

**Supplementary Table S1. VMRs defined in neurons, glia, B cells, T cells and fibroblasts**.

**Supplementary Table S2. Enrichment analysis for various genomic features overlapping VMRs in five cell types**.

**Supplementary Table S3. Networks of co-regulated VMRs defined by WCGNA in neurons, glia, B cells, T cells and fibroblasts**.

**Supplementary Table S4. Genes associated with networks of co-regulated VMRs defined by WCGNA in neurons, glia, B cells, T cells and fibroblasts**. Based on the Networks of co-regulated VMRs defined by WCGNA (Supplementary Table S3), VMRs were associated with the genes they regulate based on either their localization within ±2kb of transcription start sites, overlap with DNAseI hypersensitive sites that showed significant association *in cis* with gene expression levels (Sheffield et al., 2013), and significant associations between methylation and gene expression levels (eQTMs) in T cells, B cells, fibroblasts (Gutierrez-Arcelus et al., 2013) and monocytes (Liu et al., 2013).

**Supplementary Table S5. Significantly enriched Gene Ontology (GO) categories associated with genes linked with networks of co-regulated VMRs in neurons, glia, B cells, T cells and fibroblasts**. For each module with at least 10 constituent genes, we performed Gene Ontology enrichment analysis using GOrilla (Eden et al., 2009).

**Supplementary Table S6. Significantly enriched transcription factor binding sites overlapping networks of co-regulated VMRs defined by WCGNA in neurons, glia, B cells, T cells and fibroblasts**.

**Supplementary Table S7. VMRs defined by analysis of methylation in 26 individuals, representing 13 pairs of monozygotic twins**.

**Supplementary Table S8. VMRs identified in cultured isogenic fibroblasts grown under conditions of increasing cell density and nutrient deprivation**.

**Supplementary Table S9. Results of GO enrichment analysis using genes associated with VMRs identified in cultured isogenic fibroblasts grown under conditions of increasing cell density and nutrient deprivation**.

